# Acetylation of isoniazid - a novel mechanism of isoniazid resistance in *Mycobacterium tuberculosis*

**DOI:** 10.1101/2020.02.10.941252

**Authors:** K. B. Arun, Aravind Madhavan, Billu Abraham, M. Balaji, K. C. Sivakumar, P. Nisha, R. Ajay Kumar

## Abstract

Isoniazid (INH), one of the first-line drugs used for the treatment of tuberculosis, is a pro-drug which is converted into its active form by the intracellular KatG enzyme of *Mycobacterium tuberculosis*. The activated drug hinders cell wall biosynthesis by inhibiting InhA protein. INH resistant strains of *M. tuberculosis* usually have mutations in *katG*, *inhA*, *ahpC*, *kasA*, and *ndh* genes. However, INH resistant strains which do not have mutations in any of these genes are reported, suggesting that these strains may adopt some other mechanism to become resistant to INH. In the present study we characterized Rv2170, a putative acetyltransferase in *M. tuberculosis*, to elucidate its role in inactivating isoniazid. The purified recombinant protein was able to catalyze transfer of acetyl group to INH from acetyl CoA. HPLC and LC-MS analyses showed that following acetylation by Rv2170, INH is broken down into isonicotinic acid and acetylhydrazine. Drug susceptibility assay and confocal analysis showed that *M. smegmatis*, which is susceptible to INH, is not inhibited by INH acetylated with Rv2170. Recombinant *M. smegmatis* and *M. tuberculosis* H37Ra overexpressing Rv2170 were found to be resistant to INH at minimum inhibitory concentrations that inhibited wildtype strains. In addition, intracellular *M. tuberculosis* H37Ra overexpressing Rv2170 survived better in macrophages when treated with INH. Our results strongly indicate that Rv2170 acetylates INH, and this could be one of the strategies adopted by at least some *M. tuberculosis* strains to overcome INH toxicity.

---

For centuries tuberculosis (TB), caused by *Mycobacterium tuberculosis* (MTB), has been one of the top reasons for death of humans. WHO reports that 1.3 million people died due to TB in 2017 (1). TB is generally treated with first-line drugs isoniazid, rifampicin, ethambutol, and pyrazinamide (2). Infection by MTB resistant to at least isoniazid and rifampicin leads to multidrug-resistant (MDR) TB. Treatment of MDR-TB is more difficult and costlier than treatment of drug-susceptible TB, and the drugs cause severe side effects in patients.

Isoniazid (INH) is a potent anti-TB molecule and the killing of MTB is triggered when KatG enzyme of MTB activates the pro-drug INH by coupling it with NADH. The activated INH inhibits InhA protein by binding to its active site which in turn hinders mycolic acid synthesis, thereby disturbing the cell wall biosynthesis of MTB (3). Mutations at codon 315 of *katG* and the promoter region of *inhA* genes are often observed in INH resistant MTB. The extent of mutation in *katG* (~72%) and *inhA* (~18%) strongly depends on the genomic region as well as on the ethnic background of the patient (4). INH resistant strains with mutations in genes other than *katG* and *inhA* have also been identified and these genes include *ahpC*, *kasA* and *ndh* (5). Interestingly, about 10% of INH resistant strains do not have mutations in any of these genes (6), which suggests that these strains may have mutations in other genes or they may adopt some other strategies to counter INH.

Mechanisms by which drug resistant pathogens achieve antimicrobial resistance include degradation or modification of drugs, modification of the targets of drugs, and altering the permeability of bacterial membrane towards those drugs (7). Drug resistant pathogens often possess specialized enzymes which can modify or degrade drugs (8). Acetylation is one of the well-known mechanisms adopted by bacteria to modify drugs as well as drug targets (9). In our laboratory we have isolated an acetyltransferase, Rv3423.1, from MTB H37Rv strain which was shown to acetylate histone H3 (10). Subsequent analysis of MTB genome using the MycoBrowser portal (https://mycobrowser.epfl.ch/) revealed the presence of 11 more genes capable of coding for acetyltransferases. Bacterial proteins under GCN5-related N-acetyltransferases family are shown to be involved in drug resistance (11). We cloned two of the putative acetyltransferases (Rv0428c and Rv2170) with N-acetyltransferase domain and checked the histone acetyltransferase activity of the purified enzymes. However, we could not find any histone acetyltransferase activity for these proteins. At this point we wondered if they could be involved in the acetylation of other substrates. Many of the drugs used in the treatment of human diseases are known to be acetylated by liver enzymes. Isoniazid (INH), a first-line anti-TB drug, is known to be acetylated in the liver and gastrointestinal tract of patients to form N-acetyl INH which is further broken down into isonicotinic acid and acetylhydrazine (12). We hypothesized that MTB may employ a similar mechanism, at least in some cases, to detoxify INH. To test our proposition we checked the ability of the purified enzymes to acetylate INH *in vitro*.

## Results

### Purified Rv2170 is able to acetylate isoniazid

Rv2170 and Rv0428c were cloned into pET32a vector and heterologously expressed in *E. coli* Rosetta BL21(DE3) strain. As the recombinant proteins were His-tagged, purification was carried out by affinity chromatography using Ni-NTA agarose resin. The eluted protein fractions were checked and purity was confirmed by SDS-PAGE. Highly purified Rv2170 (23k Da) and Rv0428c (33 kDa) were observed in the gel. We confirmed the identity of the purified proteins by western blot using His tag monoclonal antibody (Fig 1A and 1B). The acetyltransferase activity of these putative acetyltransferases was checked by fluorometric assay using INH as substrate. Rv2170 exhibited significant acetyltransferase activity (Fig 1C), whereas Rv0428c did not show any activity, and therefore Rv2170 was selected for further studies.

**Figure 1.**
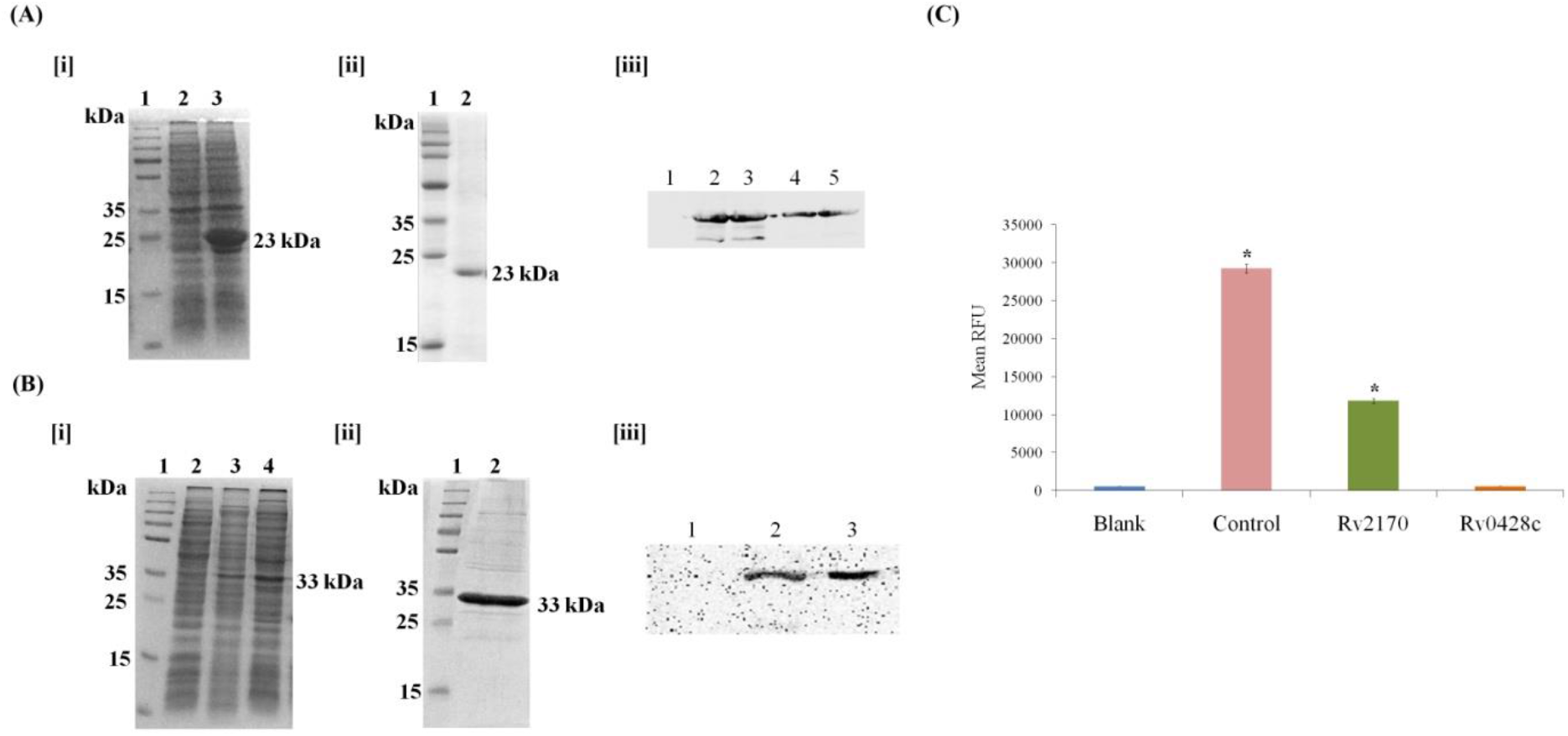
Comparison of acetyltransferase activity of two putative GNAT proteins of *M. tuberculosis*. (A) [i] *Rv2170* cloned in pET32a vector and overexpressed in *E. coli* Rosetta BL21(DE3). Lane 1-Protein Ladder, Lane 2- Uninduced, Lane 3- Induced. [ii] Purified Rv2170. Lane 1- Protein Ladder, Lane 2- Purified Rv2170. [iii] Confirmation of identity of the recomibinant Rv2170 by western blot using His-tag antibodies. Lane 1- Uninduced, Lanes 2&3- Induced, Lanes 4&5- Purified Rv2170. (B) [i] *Rv0423c* cloned in pET32a vector and overexpressed in *E. coli* Rosetta BL21(DE3) strain. Lane 1-Protein Ladder, Lane 2- Uninduced, Lanes 3&4- Induced. [ii] Purified Rv0423c. Lane 1- Protein Ladder, Lane 2- Purified Rv0423c. [iii] Confirmation of identity of the recombinant Rv0423c by western blot using His-tag antibodies. Lane 1- Uninduced, Lane 2- Induced, Lane 3- Purified Rv0423c. (C) Comparison of acetyltransferase activity of Rv2170 and Rv0428c, the two putative acetyltransferases by fluorometric acetyltransferase activity assay using isoniazid as substrate.

### Isoniazid treated with Rv2170 protein failed to kill M. smegmatis

By REMA we observed that the minimum inhibitory concentration (MIC) of INH against *M. smegmatis* was 8 μg/mL, as reported earlier (13). To test if modified INH inhibits *M. smegmatis*, the bacteria were treated with the reaction mixture (INH acetylated with Rv2170 *in vitro*) to analyze its sensitivity towards acetylated INH. REMA results (Fig 2A) clearly indicated that *M. smegmatis* remained viable when the cells were incubated with INH treated with Rv2170 (test) whereas *M. smegmatis* treated with the control (the same concentration of INH without Rv2170) were killed. The same has been proven by LIVE/DEAD assay also. Confocal microscopy (Fig 2B) and the ratio of green to red fluorescence (Fig 2C) confirmed that Rv2170 rendered INH inactive. The ratio obtained for untreated *M. smegmatis* was 5.108 ± 0.097, whereas for the bacteria treated with INH (control) the ratio decreased to 2.405 ± 0.09. However, when *M. smegmatis* was treated with acetylated INH (test) the ratio increased to 3.865 ± 0.298, strongly suggesting that Rv2170 effectively inactivates INH and hence the bacteria retained their viability. Rifampicin was used as the positive control for both REMA (1 μg/mL) and LIVE/DEAD assays (2 μg/mL). LIVE/DEAD assay performed with the end products, isonicotinic acid and acetylhydrazine, showed that these compounds were not toxic to the bacteria (Supplementary Figure 1).

**Figure 2.**
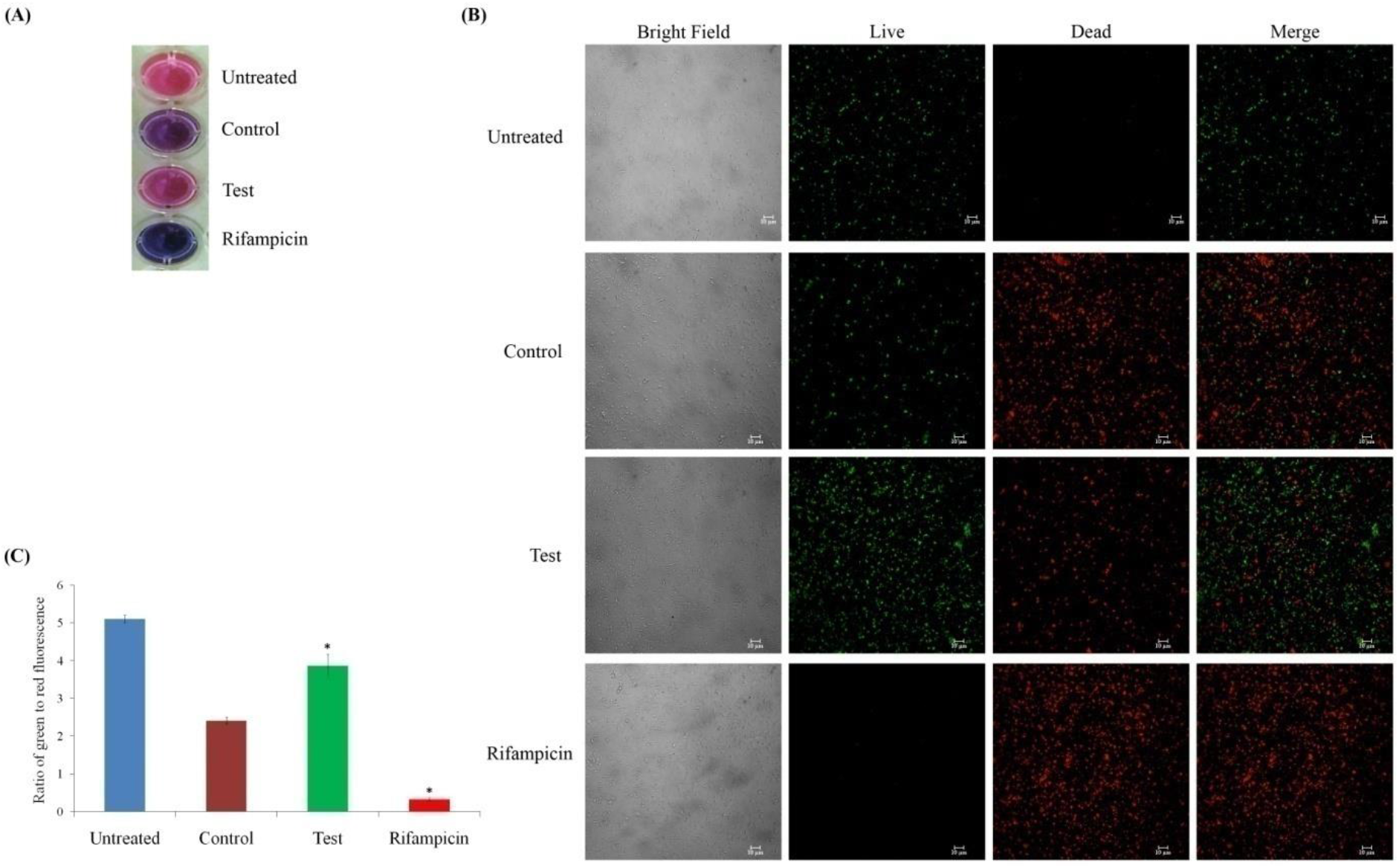
Isoniazid is inactivated by Rv2170. (A) REMA of *M. smegmatis* in the presence of INH treated with Rv2170. (B) Confocal microscopy of the *M. smegmatis* after incubating with INH treated with Rv2170. The green (SYTO 9) and red (Propidium Iodide) fluorescence show live and dead bacteria, respectively. Scale bar - 10 μm. (C) Analysis of relative viability of *M. smegmatis* after incubating with INH treated with Rv2170 by measuring the ratio of green to red fluorescence. *represents the significance of difference from the control.

### Acetylated isoniazid gets broken down into isonicotinic acid and acetylhydrazine

LC-MS analysis of the reaction mixture of INH (160 μg/mL) treated with Rv2170 (500 nM) *in vitro* revealed the presence of four compounds, whereas in the control a single peak was detected (Fig 3A). The molecular masses of three peaks INH (137.13), isonicotinic acid - INA (123.11) and acetylhydrazine - AH (74.08) were confirmed by running the respective standard molecules under the same conditions. Interestingly the fourth peak represented a molecular mass of 179.20 which very closely matched with the molecular mass of the intermediate compound acetylisoniazid - AcINH (C_8_H_9_N_3_O_2_ - 179.18). We performed HPLC analysis and the peaks in chromatograms of control and reaction mixture were identified by comparing the retention time of authentic standard molecules (INH–2.492 min, INA–3.208 min, AH–3.610 min) and confirmed by spiking the reaction mixture with them (Fig 3B and 3C). Thus these results support our hypothesis that Rv2170 acetylates INH, and the product gets broken down into INA and AH.

**Figue 3.**
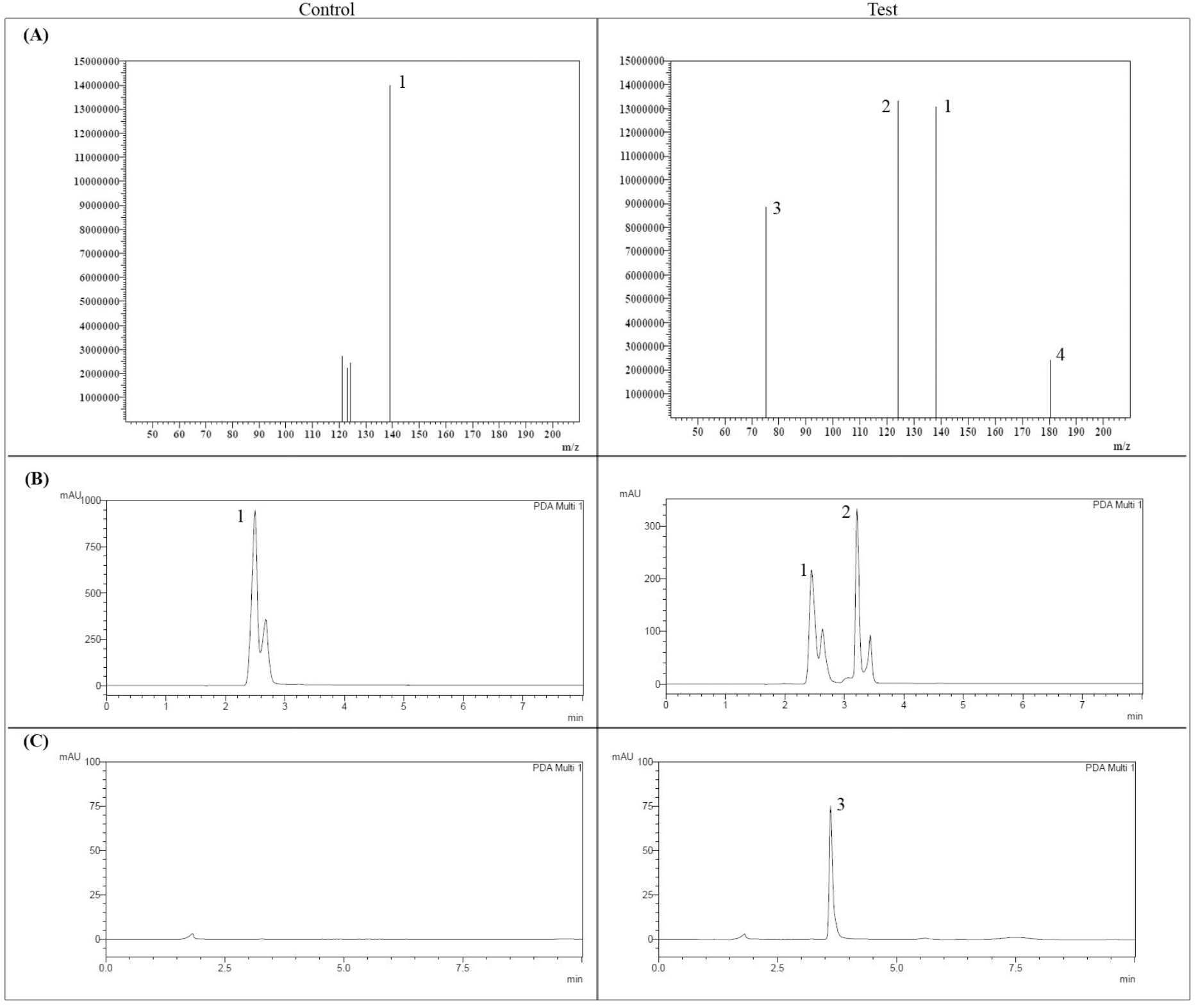
Isoniazid is acetylated into acetylisoniazid by Rv2170. Acetylisoniazid is broken own into isonicotinic acid and acetylhydrazine. (A) LC-MS analysis of *in vitro* acetylation of INH by Rv2170. Mass spectrum scanning detected four peaks in test sample – (1) isoniazid (137.13), (2) isonicotinic acid (123.11) and (3) acetylhydrazine (74.08) were confirmed with authentic molecules. The peak (4) with m/z 179.20 is almost equal to the molecular mass of the acetylisoniazid (C_8_H_9_N_3_O_2_ - 179.18), the product formed by the acetylation of isoniazid by Rv2170. (B) & (C]) HPLC analysis of *in vitro* acetylation of isoniazid (1) by Rv2170 confirms the formation of isonicotinic acid (2) and acetylhydrazine (3) in test samples.

### MSMEG_4238 - the orthologous protein of Rv2170 in M. smegmatis is less efficient in acetylating isoniazid

MSMEG_4238 is the orthologous protein of Rv2170 in *M. smegmatis*. We heterologously expressed the gene in *E. coli* and purified MSMEG_4238 and analysed its acetylation efficacy in comparison with that of Rv2170. The *in vitro* acetylation reaction mixture [INH (16 μg/mL) + 50 nM of Rv2170 or MSEMG_4238] was subjected to REMA [Fig 4(A)], LIVE/DEAD assay [Fig 4(B and C)] and HPLC analysis [Supplementary Fig 2]. The ratio obtained for untreated *M. smegmatis* was 5.096 ± 0.0781, whereas for *M. smegmatis* treated with INH (control) the ratio decreased to 2.536 ± 0.185. However, when *M. smegmatis* was treated with INH acetylated with Rv2170 (test 1) the ratio increased to 4.029 ± 0.109, and that for INH acetylated with MSEMG_4238 (test 2) increases marginally to 2.872 ± 0.157 showing that MSEG_4238 is less effective than Rv2170 protein in acetylating INH. We quantified the extent of acetylation of INH from the area of the peaks in the HPLC chromatograms [Fig 4(D)]. When Rv2170 was used for acetylation, the INH concentration declined to 4.004 ± 0.243 μg/mL from 15.302 ± 0.410 μg/mL (control), whereas for MSMEG_4238 the INH concentration came down only to 11.486 ± 1.310 μg/mL. The better activity of Rv2170 than of MSMEG_4238 is reflected in the formation of the degradation products also. The quantification results showed that acetylation of INH catalysed by Rv2170 leads to the production of more INA and AH (6.815 ± 0.027 and 1.547 ± 0.017 μg/mL, respectively), which was significantly high when compared to the activity of MSMEG_4238 (INA - 4.058 ± 0.257 μg/mL and AH - 0.340 ± 0.01 μg/mL). Thus the results obtained from these data clearly show that MSMEG_4238 is less effective than Rv2170 in acetylating INH.

**Figure 4.**
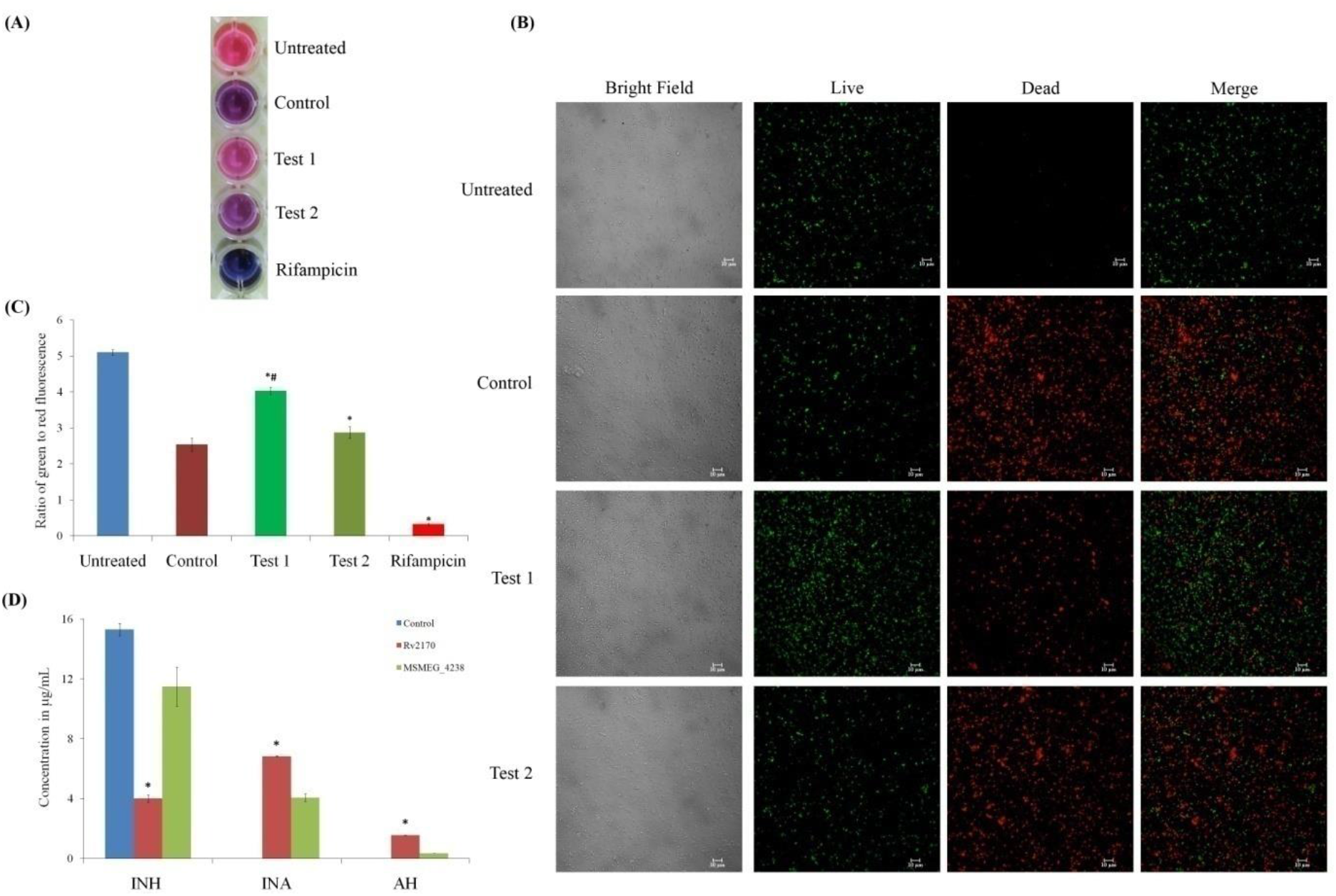
Rv2170 is more efficient in acetylating isoniazid than its orthologous protein MSMEG_4238 from *M. smegmatis*. (A) REMA of *M. smegmatis* in the presence of INH treated with Rv2170 (Test 1) and MSMEG_4238 (Test 2). (B) Confocal microscopy of the *M. smegmatis* after incubating with INH treated with Rv2170 (Test 1]) and MSMEG_4238 (Test 2). The live and dead bacteria are stained green and red respectively. Scale bar - 10 μm. (C) Analysis of relative viability of *M. smegmatis* after incubating with INH treated with Rv2170 [Test 1] and MSMEG_4238 [Test 2] by measuring the ratio of green to red fluorescence. *significantly different from control sample, p≤0.05. ^#^Test 1 significantly different from Test 2, p≤0.05. (D) Quantification of isoniazid (INH), isonicotinic acid (INA) and acetylhydrazine (AH) after *in vitro* acetylation of INH by Rv2170 based on HPLC analysis. *Rv2170 significantly different from control and MSMEG_4238, p≤0.05

### INH binds better to Rv2170 than to MSMEG_4238

We aligned the amino acid sequences of Rv2170 and MSMEG_4238 proteins using CLUSTAL O (1.2.4) and found that there are substitutions of 30 amino acids in the latter [Fig. 5(A)]. Change in structure due to the differences in amino acids may have affected the acetylation efficacy of this enzyme. BLAST search yielded the most appropriate template (PBD 1U6M) of the crystal structure of acetyltransferase with a resolution of 2.4 Å which served as the best template for building the 3D structure of Rv2170 and MSMEG_4238. They were modelled using SWISS MODEL using the template - 1u6m.1.A acetyltransferase of GNAT family from *Enterococcus faecalis*. The best-docked conformation showed that INH interacts with Rv2170 and MSMEG_4238 through hydrogen bond interactions and the drug is entirely buried in the interior of the binding pocket with its heterocyclic ring pointing towards the outer region [Figure 5 (B & C)]. The binding energy clearly showed that INH has better affinity towards Rv2170 (−5.329 kcal/mol) than to MSMEG_4238 (−4.593 kcal/mol).

**Figure 5.**
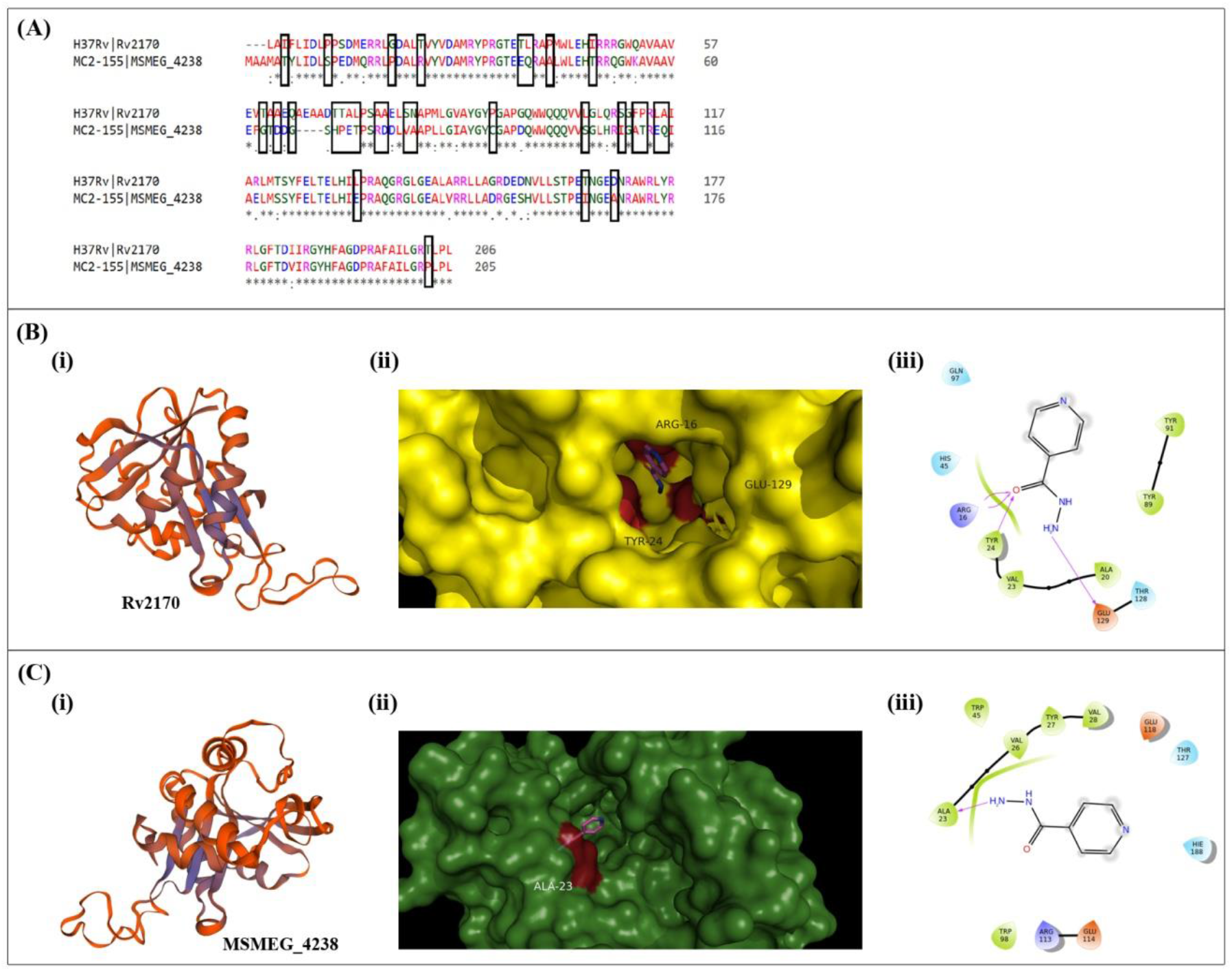
Homology modelling of Rv2170 and MSMEG_4238, and docking studies with isoniazid. (A) Sequence alignment of Rv2170 and its orthologous protein MSMEG_4238. Amino acid substitutions are indicated in boxes. (B) Homology model of Rv2170 (i), and superimposed image (ii) and schematic representation of interaction with INH (iii). (C) Homology model of MSMEG_4238 (i), and superimposed image (ii) and schematic representation of interaction with INH (iii).

### M. smegmatis and M. tuberculosis H37Ra overexpressing Rv2170 are resistant to INH

The MIC values for INH against *M. smegmatis* and MTB were determined to be 8 and 0.5 μg/mL, respectively (Supplementary figure 3 and 4) which matched with the concentrations reported earlier (14, 15). To assess whether the enzyme offers any survival advantage to *M. smegmatis* and MTB H37Ra in the presence of INH, we overexpressed *Rv2170* in these bacteria. The recombinant strains were exposed to INH (final concentration 8 and 0.5 μg/mL, for *M. smegmatis* and MTB H37Ra, respectively) and then performed REMA, LIVE/DEAD assay and CFU count. Bacteria carrying empty pBEN vector served as control. REMA results clearly showed that the recombinant *M. smegmatis* survived INH (8 μg/mL) toxicity whereas *M. smegmatis* carrying empty vector succumbed to the drug toxicity (Fig 6A). This was also evident from fluorescent data (Fig 6B) and LIVE/DEAD confocal analysis (Fig 6C). The green to red ratio obtained for *M. smegmatis* (vector control) treated with INH was 2.468 ± 0.091, and 3.868 ± 0.047 for recombinant *M. smegmatis* suggesting that *M. smegmatis* overexpressing *Rv2170* can resist INH toxicity.

**Figure 6.**
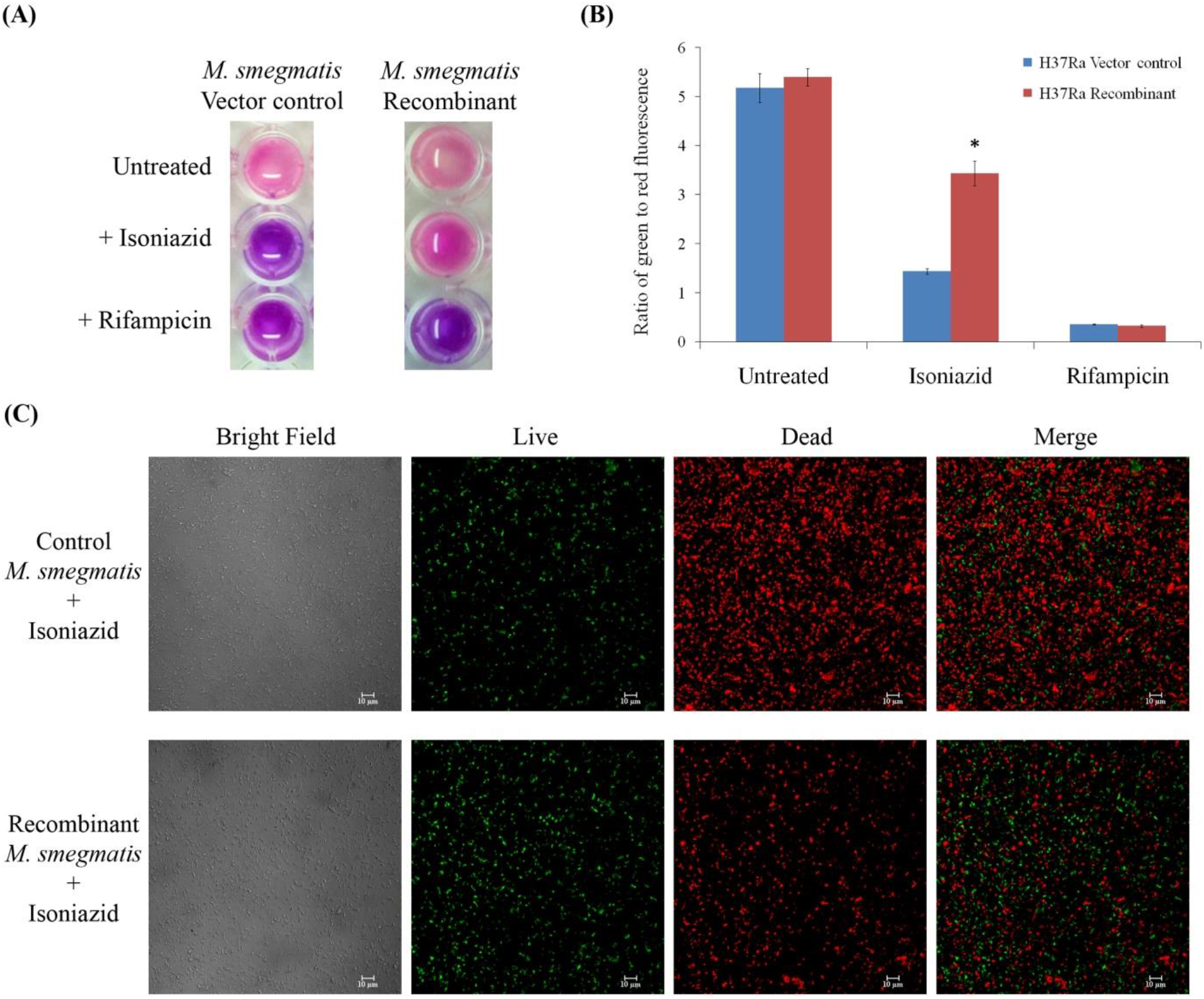
Recombinant *M. smegmatis* overexpressing *Rv2170* is resistant to INH. (A) REMA of *M. smegmatis* (vector control and recombinant) in the presence of INH. (B) The relative viability of vector control and recombinant *M. smegmatis* after incubating with INH, respectively, by measuring the ratio of green to red fluorescence. (C) Confocal microscopy of *M. smegmatis* (vector control) and recombinant *M. smegmatis* carrying *Rv2170* after incubating with INH. The green (SYTO 9) and red (Propidium Iodide) fluorescence show live and dead bacteria, respectively. Scale bar - 10 μm.

Recombinant MTB H37Ra strain expressing *Rv2170* retained its viability after treating with INH whereas the vector-alone control did not. REMA results (Fig 7A) showed that the recombinant MTB effectively resisted INH (0.5 μg/mL) when compared to the vector-alone control. Green to red fluorescence ratio (Fig 7B) for MTB (vector control) treated with INH was 1.436 ± 0.056 and it was 3.434 ± 0.252 for the recombinant bacterium, which clearly showed that MTB that overexpressed *Rv2170* could resist INH toxicity. Results from CFU assay also showed that MTB overexpressing *Rv2170* could effectively survive INH toxicity (Fig 7C). When MTB (vector control) was treated with INH the log CFU obtained was 3.611 ± 0.038 while the number increased significantly (5.316 ± 0.270) when recombinant MTB was treated with INH.

**Figure 7.**
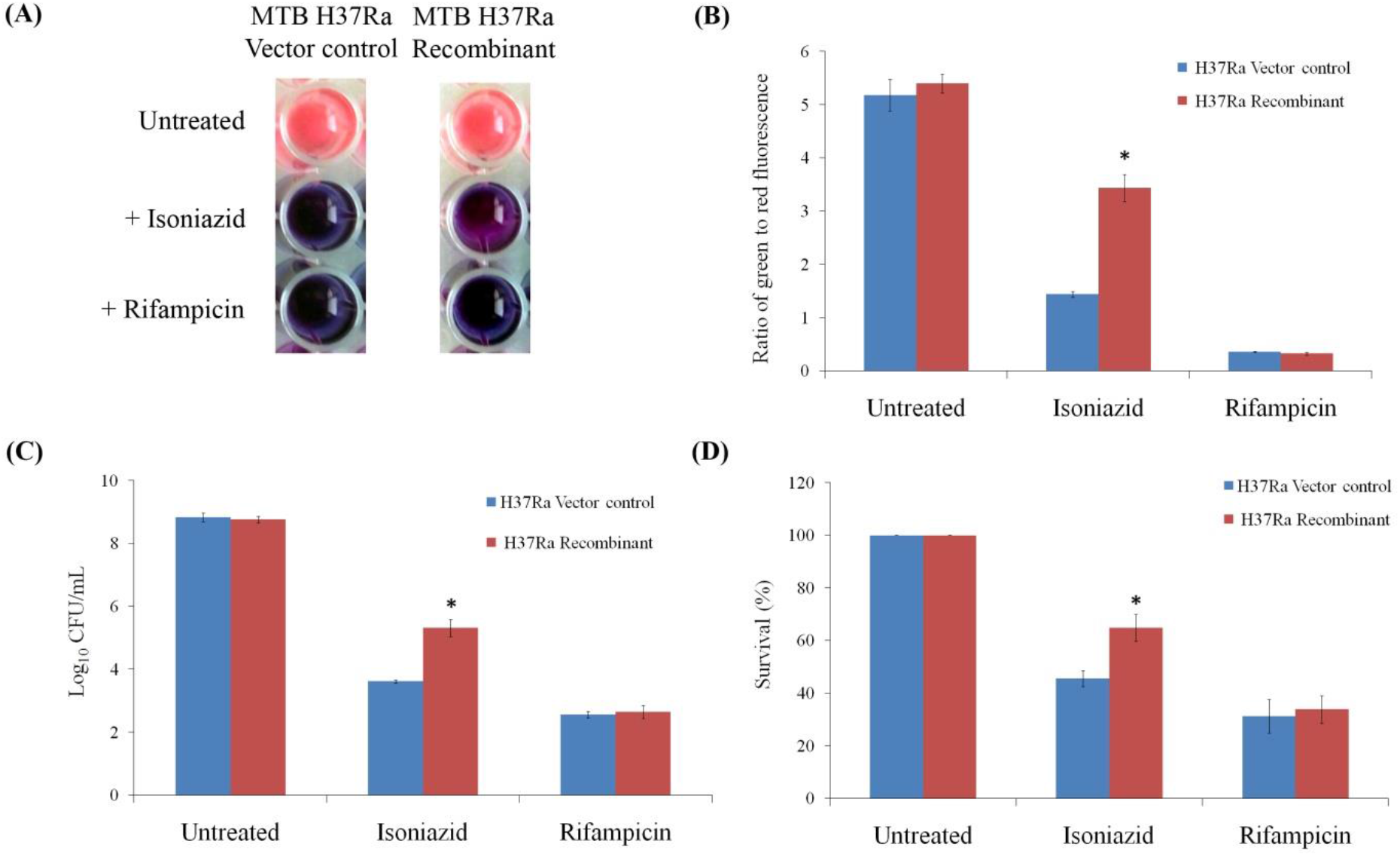
Recombinant *M. tuberculosis* H37Ra overexpressing *Rv2170* survives better in macrophage when treated with INH. (A) REMA of H37Ra (vector control and recombinant) in the presence of INH. (B) Analysis of the ratio of green to red fluorescence to measure the relative viability of vector control and recombinant H37Ra after incubating with INH. (C) Measurement of bacterial viability by CFU count. The graph represents the log value of CFU obtained after treating with INH. (D) The intracellular survival of recombinant *M. tuberculosis* H37Ra in macrophages after treating with INH. *recombinant group significantly different from vector control group, p≤0.05.

### M. tuberculosis H37Ra overexpressing Rv2170 showed better intracellular survival in macrophages treated with INH

To analyse whether the protein offers any protective advantage to the intracellular bacteria in the presence of INH, THP-1 cells infected with MTB H37Ra overexpressing *Rv2170* or carrying empty vector were treated with INH for 24 h. The CFU counts obtained revealed that INH fails to kill intracellular recombinant MTB overexpressing *Rv2170*. The survival percentage (Fig 7D) of recombinant MTB (64.82 ± 5.05%) was significantly higher than that of MTB vector-alone control (45.54 ± 3.09%) suggesting that overexpression of *Rv2170* in MTB enhances its survival in macrophages in the presence of INH.

## Discussion

Multidrug resistant strains of *M. tuberculosis* tolerate the key first-line drugs isoniazid and rifampicin and this has negatively impacted the tuberculosis control programs globally (16). According to WHO report the average of isoniazid resistance is 7.2% in new TB cases, and 11.6% in previously treated TB cases (17). INH resistance mainly arises due to mutations in target genes *katG, inhA, ahpC, kasA* and *ndh* (18). In humans INH is readily absorbed by the gastrointestinal tract and is metabolized by N-acetyltransferases (NAT 2) present in liver and gastrointestinal tract into isonicotinic acid (INA) and acetylhydrazine (AH) (19). Many laboratories have reported INH resistant MTB strains with no known mutations (20, 21, 22) suggesting possible existence of other strategies to survive in the presence of INH. This made us wonder if acetyltransferases in MTB, like NAT2 of liver and gastrointestinal tract, can also inactivate INH by acetylation.

Acetylation is one of the widely adopted strategies of pathogens to deactivate drugs such as aminoglycosides, chloramphenicol, streptogramins, and fluoroquinolones (23). Chloramphenicol acetyltransferases that acetylate choramphenicol resulting in chloramphenicol resistance in bacteria have been extensively reviewed by Schwarz et al (24). *Riemerella anatipestifer* causing serositis and septicaemia in poultry are reported to show resistance towards chloramphenicol by acetylation mechanism (25). Eis protein of MTB is reported to be involved in the deactivation of capreomycin by acetylation (26). Arylamine *N*-acetyltransferase of MTB has been reported to acetylate *p*-aminosalicylic acid leading to its inactivation (27).

We screened two probable acetyltransferases of MTB (Rv0428c and Rv2170) with N-acetyltransferase domain for their ability to acetylate INH. A proprietary fluorometric assay was performed to analyse the transfer of acetyl group of acetyl CoA to INH, and the CoA released during the reaction is converted into a fluorescent product quantity of which indirectly indicates the acetylation activity. The results clearly showed that Rv2170 has the ability to acetylate INH.

Since liquid chromatography coupled with mass spectrometry is widely used in the identification of drugs and their degradation products (28), we employed the same to check and identify the metabolites formed after the treatment of INH with Rv2170 protein. LC-MS of *in vitro* acetylated mixture showed four peaks of which three complied with the molecular masses of standard isoniazid (INH), isonicotinic acid (INA) and acetylhydrazine (AH). The fourth peak obtained in LC-MS with m/z 179.20 very closely matched with the molecular mass of acetylisoniazid (AcINH, C_8_H_9_N_3_O_2_ - 179.18). AcINH is the intermediate compound formed after the acetylation of INH by Rv2170 which is broken down into INA and AH. The same has been confirmed by HPLC also, after analyzing the reaction mixture in which INH was acetylated. Since AH is aliphatic in nature we have used a different solvent system for its identification. Thus LCMS and HPLC analyses prove our hypothesis that Rv2170 effectively acetylates INH nullifying the drug toxicity. The breakdown products INA and AH were found to be non-toxic to the bacterium.

INH has an MIC of 8 μg/mL against *M. smegmatis* [the surrogate organism for virulent MTB] (13, 14, 29); employing REMA we have also obtained the same MIC value. We reasoned that if INH is rendered inactive by acetylation *in vitro*, treatment of the bacterium with this modified INH would not affect the viability of the bacteria at this concentration. *M. smegmatis* was treated with *in vitro* acetylation mixture of INH (INH + Rv2170 protein) and we subjected the treated bacteria to REMA and confocal analysis. The results showed that the bacteria retained viability when they were treated with the acetylation reaction mixture that contained INH (8 μg/mL), and the data obtained from both the experiments proved that *M. smegmatis* remained viable in the presence of INH acetylated *in vitro* by Rv2170 protein.

MSMEG_4238 is the orthologous protein of Rv2170 in *M. smegmatis* and we checked if it would also acetylate INH. Results from REMA, LIVE/DEAD assay and HPLC analysis indicated that MSMEG_4238 does possess acetyltransferase activity albeit to a lesser extent compared to that of Rv2170. However, bioinformatic analysis revealed that Rv2170 has significant similarity with its orthologous proteins present in pathogenic mycobacteria such as *M. leprae*, *M. marinum*, *M. africanum*, *M. canettii* and *M. bovis*; whereas a number of substitutions were observed in the proteins in non-pathogenic mycobacteria such as *M. smegmatis*, *M. vaccae*, *M. aurum* and *M. gilvum*. While trying to correlate their evolutionary relationship orthologs of Rv2170 from pathogenic and non-pathogenic mycobacteria formed separate clades (Supplementary Figure 5 and 6). This suggests a possibility that Rv2170 and its orthologous proteins in pathogenic mycobacteria at some point of time evolved into a virulence factor, and the pathogenic bacteria became a separate group during evolution. The ScanProsite analysis (30) of functional domains showed that amino acids from 44-205 and 31-204 form the GNAT conserved domain for Rv2170 and MSMEG_4238, respectively. Alignment of amino acids of Rv2170 and MSMEG_4238 showed 30 non-conserved substitutions in the conserved domain; for example Leu105 and Thr203 of Rv2170 are substituted by Ser and Pro in MSMEG_4238, respectively. The difference in amino acids may have altered the protein conformation of MSEMG_4238 which most likely resulted in the reduced affinity of INH for the protein.

We carried out docking analysis to verify the binding efficiencies of INH to Rv2170 and MSMEG_4238. As the number of hydrogen bonds is an important factor influencing protein stability and function, hydrogen bond occupancies at a distance cut-off of 3.5 Å and an angle cut-off of 30° were applied in the calculation of hydrogen bonds. The −NH_2_ and C=O functional groups of isoniazid form strong hydrogen bond interactions with the binding site residues of the protein. In the Rv2170-INH complex, the NH_2_ moiety of INH forms hydrogen bonds with the side chain of Glu129, while C=O species makes hydrogen bonds with Arg16 and Tyr24 deep in the INH binding pocket. Meanwhile, the best binding pose of INH suggests that it is capable of forming only a single hydrogen bond between −NH_2_ moiety and Ala23 in MSMEG_4238-INH complex. The increased stability of Rv2170-INH could be due to the formation of hydrogen bonds and extensive non-polar contacts which stabilised the protein structure. Interestingly, INH interacts with Glu129 which is in the acetyltransferase domain of the Rv2170; whereas no such interaction is observed in the case of MSMEG_4238. The interaction of INH with Glu129 of functional domain of Rv2170 might have facilitated the effective transfer of the acetyl group from acetyl CoA to INH.

The recombinant *M. smegmatis* and MTB H37Ra overexpressing *Rv2170* effectively withstood INH toxicity, and recombinant MTB H37Ra showed enhanced survival inside macrophages in the presence of INH.

The proposed INH acetylation reaction catalysed by Rv2170 is shown in Figure 8. However, how acetylisoniazid gets converted into INA and AH is not clear from our study. Since this is a hydrolysis reaction (AcINH + H2O → INA + AH), the reaction probably will be catalysed by any one of the hydrolase enzymes present in MTB which is yet to be identified. Taken together, our results suggest that Rv2170-mediated acetylation of INH is a novel strategy adopted by at least some of the INH resistant MTB strains.

**Figure 8.**
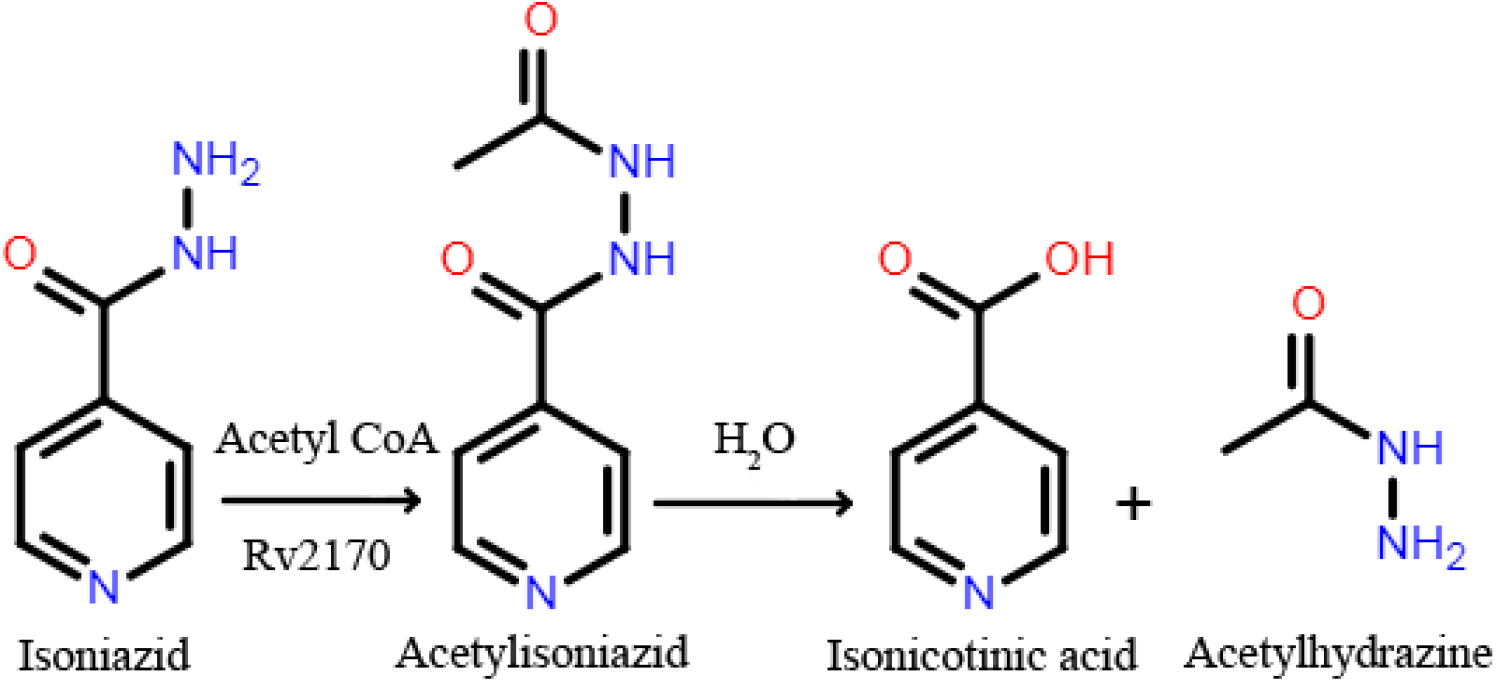
Schematic representation of acetylation of isoniazid by Rv2170, and hydrolysis of acetylisoniazid into isonicotinic acid and acetylhydrazine.

## Experimental procedures

*M. tuberculosis* H37Ra was grown in Middlebrook 7H9 medium (BD Difco™, Fisher Scientific, USA) supplemented with glycerol (0.2%), Tween 80 (0.05%) and 10% OADC (HiMedia, India). Experiments involving MTB were performed in a Biosafety Level 3 facility. *M. smegmatis* mc^2^155 and the recombinant strains were grown in LB broth supplemented with 0.05% Tween 80 or in Middlebrook 7H9 medium (Difco, Detroit, MI, USA) supplemented with 0.5% glycerol, 0.05% Tween 80 and 10% oleate/albumin/dextrose/catalase supplement (Becton Dickinson, Franklin Lakes, NJ, USA) at 37 °C. *Escherichia coli* strains JM109 (Promega, Madison, WI USA) and BL21(DE3) Rossetta, (Novagen, Madison, WI, USA) were used for cloning and expression of genes, respectively. pET32a vector (Novagen) was used for recombinant gene expression in *E. coli* and pBEN vector was used for expression in *M. smegmatis* and MTB H37Ra.

Genes of two putative acetyltransferases Rv0428c and Rv2170, with GCN5-related N-acetyltransferase domain, were amplified from the genomic DNA of MTB H37Rv that was available in the laboratory using specific primers (Supplementary Table 1). The PCR products were purified and were cloned into a pET32a vector using *Nde*1 and *Hin*dIII restriction sites. The resulting construct was then transformed into competent *E. coli* (JM109) cells. Ampicillin resistant clones were selected, and the recombinant plasmid was then transformed into competent *E. coli* BL21(DE3) Rossetta cells. Isopropyl β- D -1-thiogalactopyranoside (IPTG - 1 mM) was used to induce expression of the genes. The over-expressed 6X His-tagged fusion proteins were purified under native conditions using Ni-NTA agarose resin (Thermo Scientific, Waltham, MA, USA). The purified proteins were concentrated and buffer-exchanged using Amicon Ultra-15 centrifugal filter units (Millipore, Billerica, MA, USA) and stored in 20% glycerol at −20 °C for assays.

Acetyltransferase activity of Rv0428c and Rv2170 was determined using fluorometric acetyltransferase activity assay kit (ABCAM-ab204536, Cambridge, USA) and isoniazid (100 μM) as substrate. The assay was performed with recombinant protein (50 nM) according to manufacturer’s instructions.

The *in vitro* acetylation assay was performed according to Kuninger et al. (31) with slight modification. *In vitro* acetylation reaction mixture contained isoniazid (16 μg/mL = 116.67 μM), acetyl CoA (116.67 μM) and protein (50 nM) in acetyltransferase buffer (50 mM Tris HCl - pH 8.0 and 10% glycerol). The reaction was performed for 45 min at 37 °C, and was terminated by adding 0.05% H_2_SO_4_. This reaction mixture was used as the source of INH in the bacterial viability assay and confocal analysis. For LC-MS and HPLC analyses we increased the concentration of the components of the *in vitro* acetylation reaction mixture by tenfold for better detection.

The reaction mixture after performing the *in vitro* acetylation assay was subjected to liquid chromatography-mass spectrometry (LC-MS) analysis. The analysis was carried out on a UHPLC LC-30A (Shimadzu, Japan) associated with a Triple Quadrupole Mass Spectrometer LCMS-8045. The instrument was equipped with solvent delivery pumps (LC-30AD), degassing unit (DGU-20A5R), auto sampler (SIL-30AC), column oven (CTO-30AC), system controller (CBM-20A), triple quadrupole mass spectrometer (LCMS8045) and LabSolutions Ver. 5.86 software for data analysis. The assay mixture was filtered (0.22 μm N_66_Posidyne positively charged zeta membrane, Pall Life Sciences) and injected into the column (Shim-pack GISS, 1.9 μM C18, 2.1×150 mm, Shimadzu). Methanol (70%) containing 0.1% formic acid was used as the mobile phase for the LC analysis followed by mass analysis of the compounds via electrospray ionisation (ESI)-MS in the scanning mode.

Chromatographic analysis of the reaction mixture and the reference compounds was performed using a Prominence UFLC system (Shimadzu, Japan) equipped with system controller (LC-20AD), C18 column (Phenomenex Gemini, 250 × 4.6 mm, 5 μm), column oven (CTO-20A), a Rheodyne injector (20 μL loop) and a diode array detector (SPD-M20A). Twenty microlitre of sample filtered through 0.45 μm PTFE filter was injected into the column and the column temperature was maintained at 33 °C. The run time was set to 8 min. For the detection of isoniazid and isonicotinic acid, a mixture of 40% acetonitrile and 60% phosphate buffer (20 mM di-sodium hydrogen phosphate and pH was adjusted to 4.5 with ortho phosphoric acid) was used as the mobile phase with a flow rate of 0.8 mL/min. For the determination of acetylhydrazine HPLC grade water was used as the mobile phase with a flow rate of 1 mL/min. The sample peaks were monitored at 260 nm (isoniazid and isonicotinic acid) and 206 nm (acetylhydrazine). The compounds in the *in vitro* acetylation assay mixture were analyzed by comparing with standard compounds using the retention time, and also by spiking the samples with standards. Quantities of compounds in the samples were determined from the standard calibration curves plotted with known quantities of authentic commercial molecules. LC LabSolutions software was used for data acquisition and analysis.

Resazurin microtitre assay (REMA) was performed as detailed by Martin et al. (32). Bacterial viability was analysed by using LIVE/DEAD*^®^Bac*Light™ Bacterial viability kit (L7007 Molecular Probes, Invitrogen). The experiment was performed as per the manufacturer’s instructions. The bacterial cells with reagent mixture were incubated at 37 °C in the dark for 15 min. After incubation fluorescence intensity was measured at 485/530 nm (green) and 485/630 nm (red) and the ratio intensities of green to red was calculated. Following this the bacteria were observed under confocal microscope (Nikon A1R equipped with Nikon Eclipse Ti-S inverted fluorescent microscope).

To infer the structure of Rv2170 and its orthologous protein MSMEG_4238 from *M. smegmatis*, we performed BLASTP (33) search of the Protein Data Bank (PDB) using the NCBI server (http://www.ncbi.nlm.nih.gov) with default parameters for identifying suitable templates for homology modelling. Among the five homology models of Rv2170 and MSMEG_4238 generated by using MODELLER 9v12 (34), the best model was selected based on PROCHECK analysis (35) and the possible best active site was determined using the CASTp web server (36). The preparatory steps for docking studies were performed by utilising the software AutoDock Tools 4, and the grid box creation (37) for docking of ligands based on the predicted active site. All possible rotatable bonds and partial atomic charge distribution of the ligands were determined via Gasteiger methods, and the Kollman charges were assigned for all atoms of the receptor. In Autodock4.2 the structures were docked using the Lamarckian genetic algorithm with the docking parameters defined as follows: population size of 200, 100 docking runs, random starting position and conformation, mutation rate of 0.02, crossover rate of 0.8, translation step ranges of 2.0 Å, 2.5 million energy evaluations and local search rate of 0.06. The docking results <1.0 Å were clustered based on similarities in binding modes and affinities in each run discriminated by using their positional root-mean square deviation (RMSD), and represented with the most favourable free energy binding.

The *Rv2170* gene was amplified from MTB H37Rv genomic DNA with the forward (Rv2170PBENF) and reverse (Rv2170PBENR) primers (Supplementary Table 1) containing *Bam*HI and *Hin*dIII restriction sites, respectively. The gene was cloned into mycobacterial constitutive expression vector pBEN using the above-mentioned restriction sites in the multiple cloning site downstream of the mycobacterial *hsp60* promoter. Electroporation of *M. smegmatis* mc^2^155 and MTB H37Ra competent cells was performed using a Gene Pulser electroporation system (Bio-Rad) at a voltage of 1.5 kV, resistance of 200 Ω and capacitance of 25 μF. The recombinant cells were selected on Middlebrook 7H10 agar plates containing oleate-albumin-dextrose-catalase (10%) and kanamycin (25 μg/mL). MTB H37Ra (wildtype and recombinant) culture adjusted to an OD of 0.2 units at 600 nm (corresponds to 3 × 10^8^ bacteria/mL) was diluted 20 times and treated with *in vitro* acetylation mixture or rifampicin for 48 h at 37 °C. All the treated and untreated cultures were serially diluted and plated on Middlebrook 7H10 agar (BD Difco™, Fisher Scientific, USA). After incubation at 37 °C for 3 weeks the colonies were counted and colony forming units (CFU) was calculated.

THP-1 cells (20 × 10^3^) were differentiated into macrophages by inducing with phorbol 12-myristate 13-acetate (PMA - 20 ng/mL) for 24 h and were infected with wild type and recombinant H37Ra at a multiplicity of infection of 20:1. After 4 h, unphagocytosed bacteria were killed by treating with gentamycin (45 μg/mL). Subsequently the cells were treated with isoniazid (6 μg/mL) or rifampicin (1 μg/mL) for 24 h at 37 °C. The cells were lysed with sterile water and plated on Middlebrook 7H10 agar and incubated at 37 °C. The colonies were counted after 3-4 weeks and results were expressed as percentage of survival.

## Acknowledgements

Arun K B would like to acknowledge Science Engineering and Research Board (Department of Science and Technology, Government of India) and Kerala Biotechnology Commission (Kerala State Council for Science and Technology, Government of Kerala) for post-doctoral fellowship. Aravind Madhavan would like to acknowledge Department of Biotechnology for post-doctoral fellowship. The authors thank the staff of confocal microscopy facility, RGCB for their support in imaging. R Ajay Kumar thanks Department of Biotechnology, New Delhi for financial support.

## Author contributions

KBA, AM and RAK conceived and designed the experiments and interpreted the data. KBA, BA and PN performed LC-MS and HPLC and analyzed the data. KBA and AM performed confocal analysis. KBA and MB performed drug viability assays. KCS performed docking analysis. KBA wrote the manuscript and RAK critically revised the article and approved the final version to be published.

---

**Table 1.**
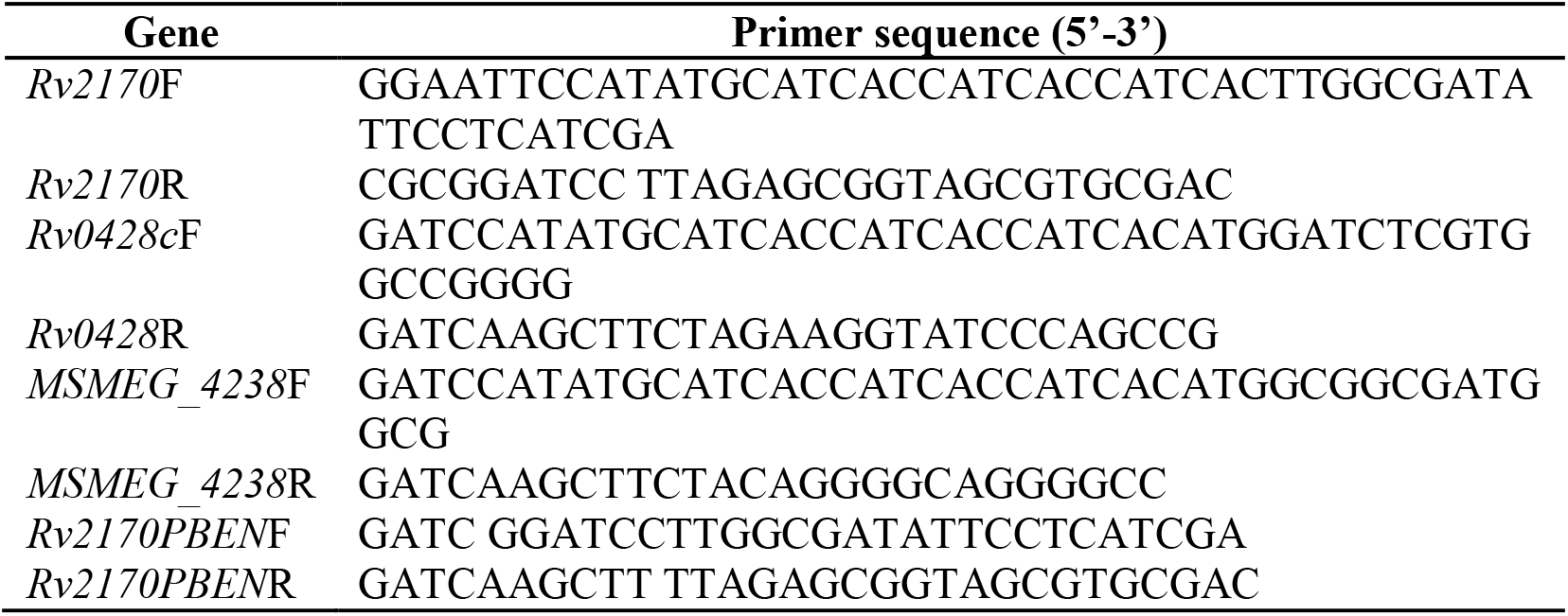
List of primers used in this study

**Supplementary Figure 1.**
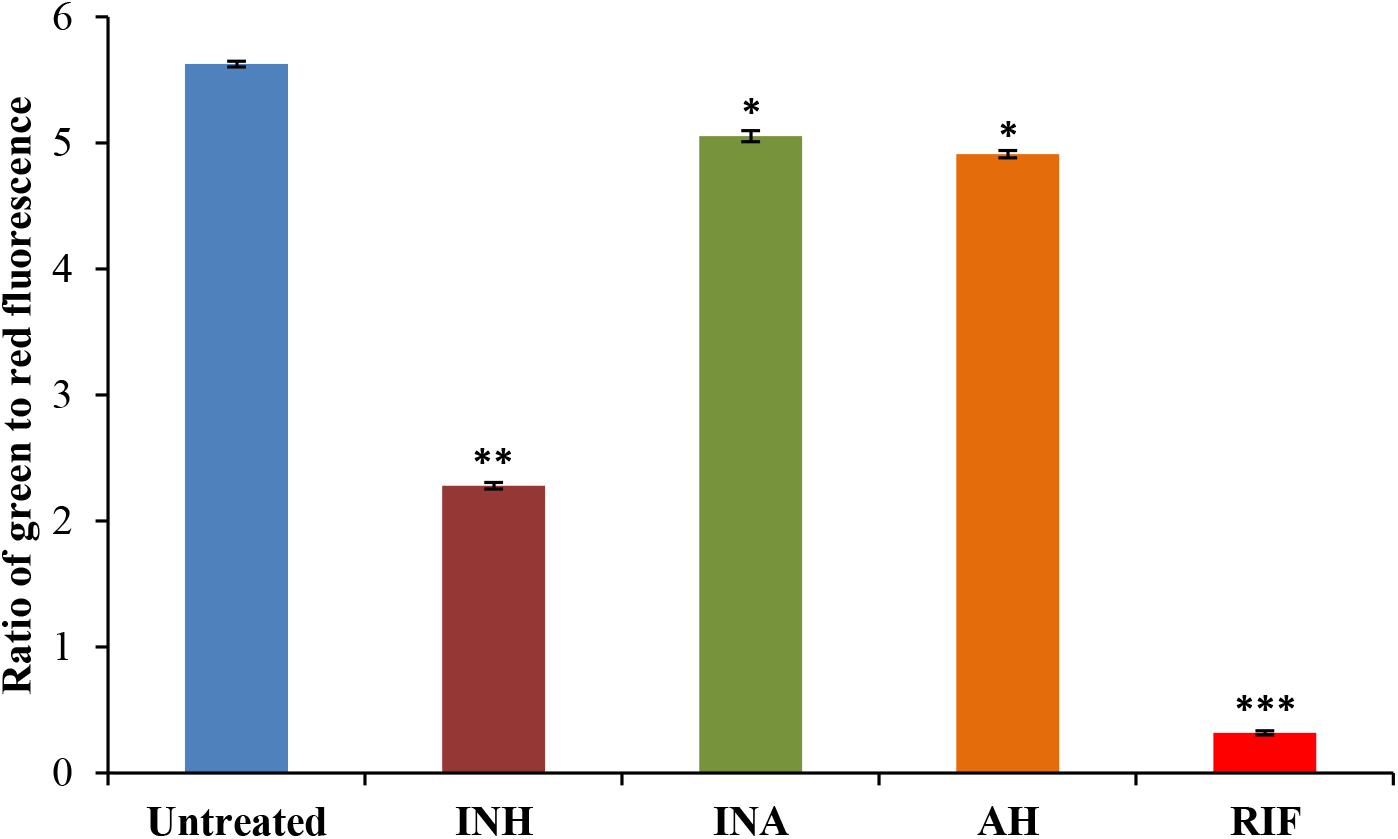
Analysis of relative viability of *M. smegmatis* after incubating with isoniazid (INH), isonicotinic acid (INA), acetylhydrazine (AH) and rifampicin (RIF) by measuring the ratio of green to red fluorescence. *significance of difference from control sample, p≤0.05

**Supplementary Figure 2.**
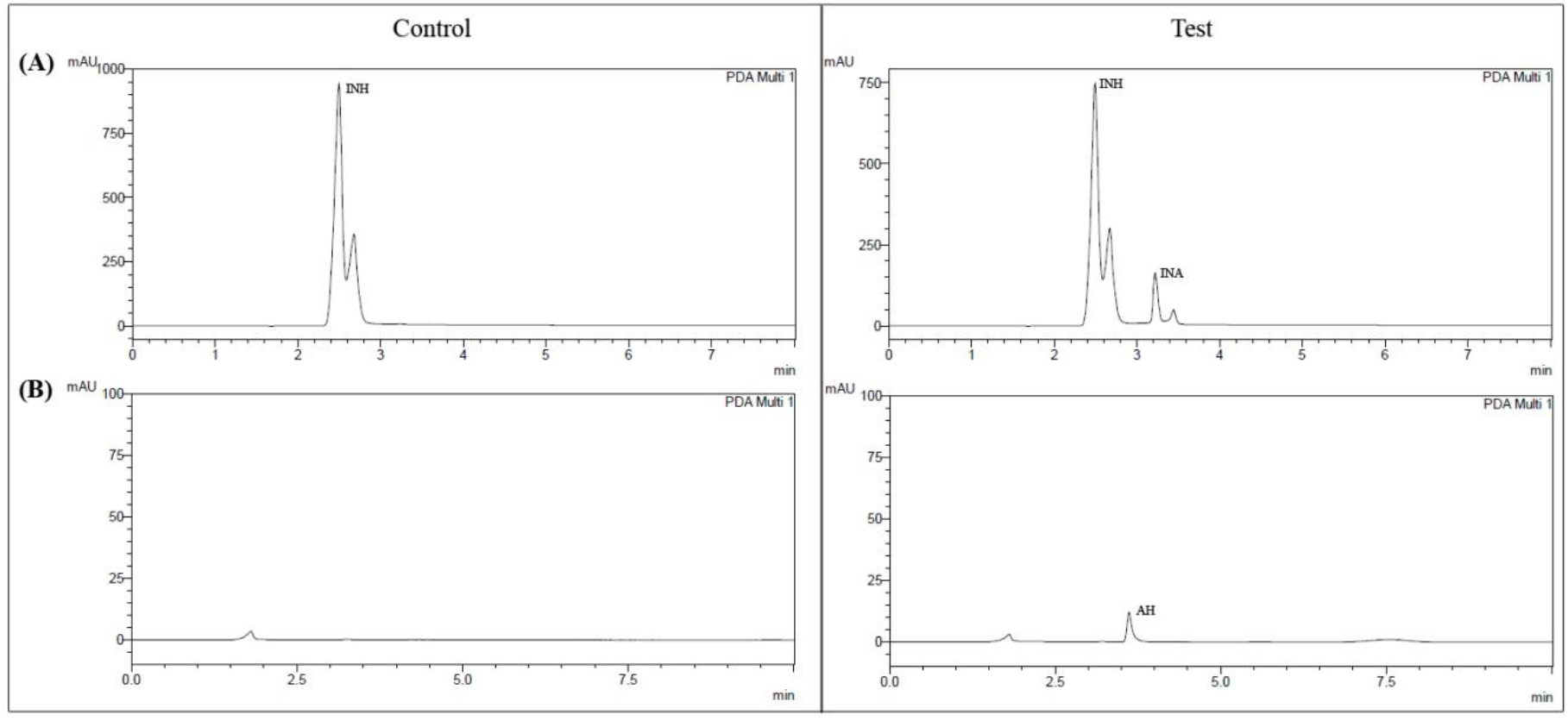
HPLC analysis of *in vitro* acetylation of isoniazid [INH] by MSMEG_4238 shows formation of very little of (A) isonicotinic acid [INA], and (B) acetylhydrazine [AH] in test samples.

**Supplementary Figure 3.**
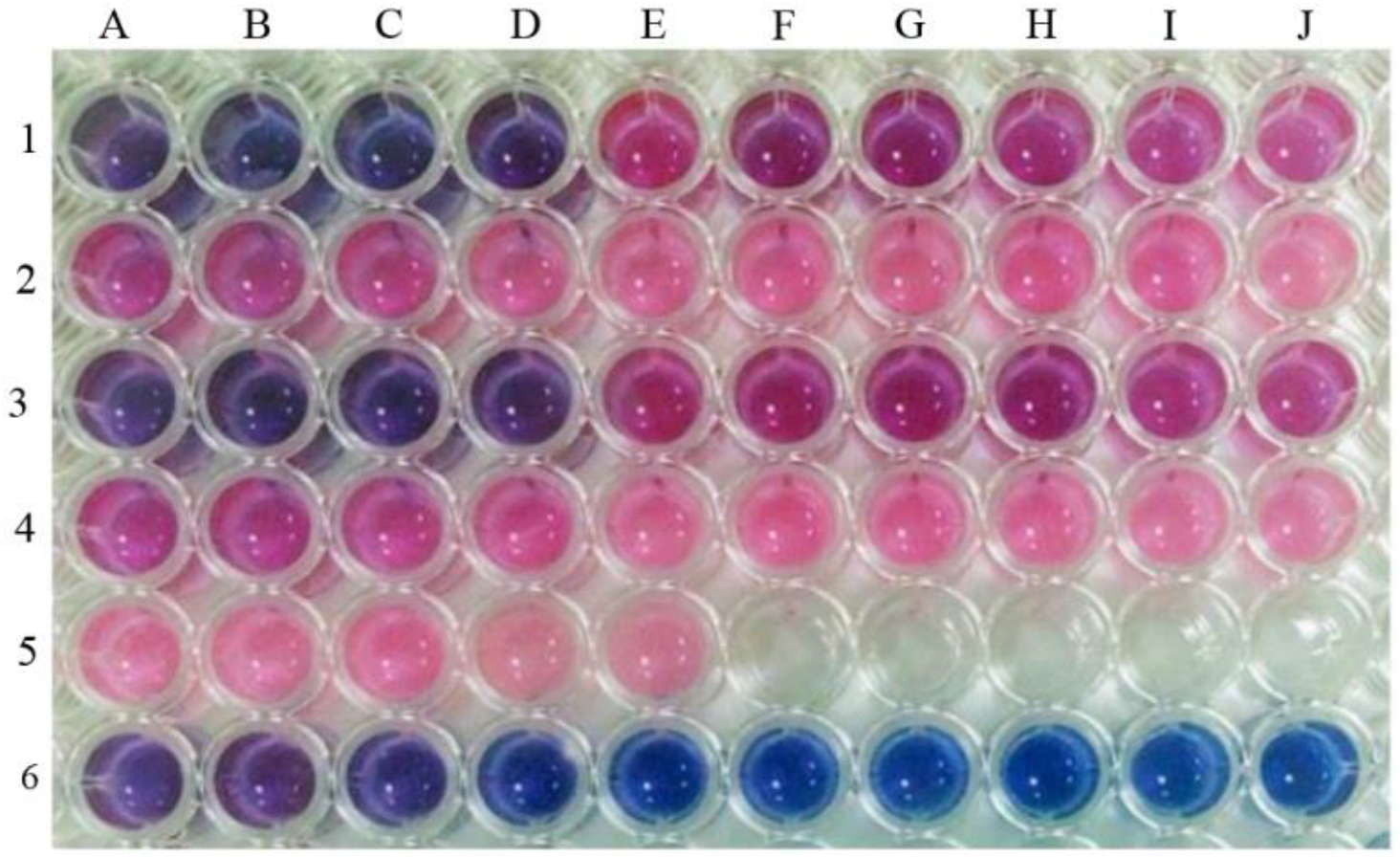
Resazurin microtiter assay (REMA) to determine the minimum inhibitory concentration (MIC) of isoniazid against *M. smegmatis*. Wells of rows 1 to 2 (1A to 2J) contain *M. smegmatis* treated with decreasing concentrations of isoniazid (64 μg/mL to 0.122 ng/mL). Wells 3A to 4J represent duplicates of the same. Wells 5A to 5E contain bacteria without any drug, and the remaining wells in row 5 contain sterile water. Wells of row 6 contain medium with isoniazid (64 μg/mL to 0.125 μg/mL) alone. Well 1D represents the MIC (8 μg/mL) of isoniazid against *M. smegmatis.*

**Supplementary Figure 4.**
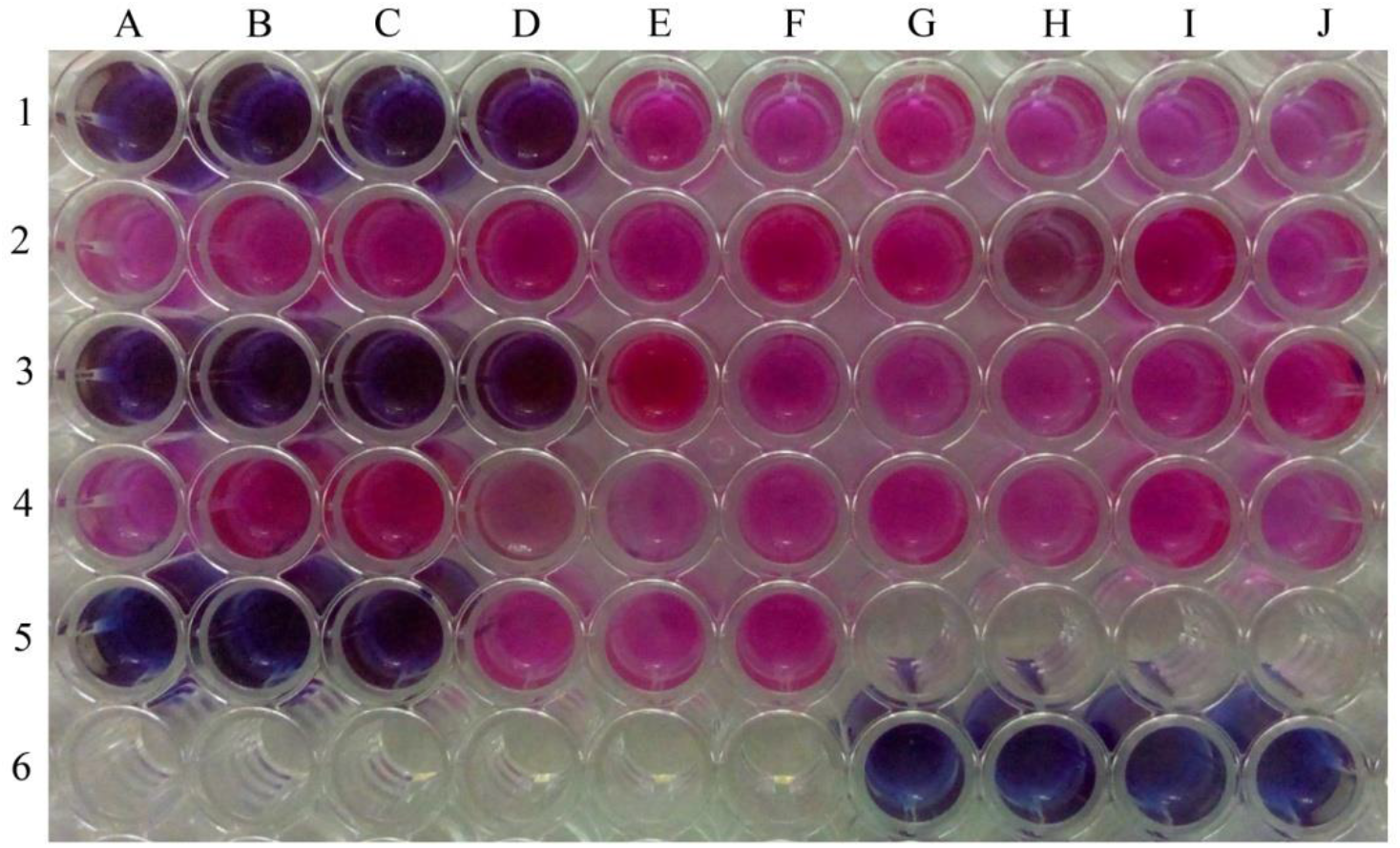
Resazurin microtiter assay (REMA) to determine the minimum inhibitory concentration (MIC) of isoniazid against *M. tuberculosis* H37Ra. Wells of rows 1 to 2 (1A to 2J) contain *M. tuberculosis* H37Ra treated with decreasing concentrations of isoniazid (4 μg/mL to 0.0076 ng/mL). Wells 3A to 4J represent duplicates of the same. Wells 5A to 5C contain bacteria treated with rifampicin (1 μg/mL). Wells 5D to 5F contain bacteria without any drug. Wells 5G to 6F contain sterile water. Wells 6G to 6J contain medium with isoniazid (4 to 0.5 μg/mL) alone. Well 1D represents the MIC (0.5 μg/mL) of isoniazid against *M. tuberculosis* H37Ra.

**Supplementary Figure 5.**
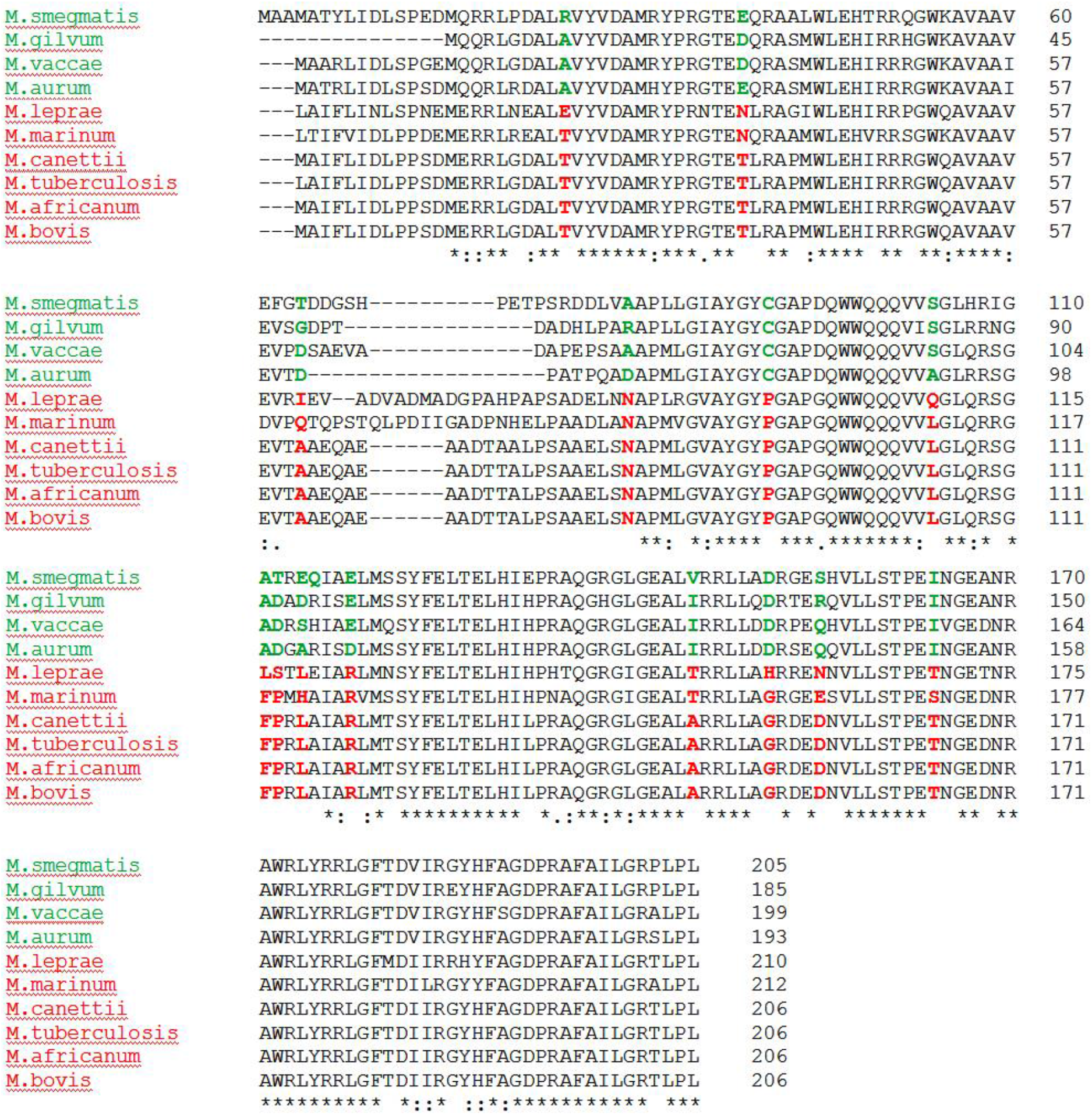
Amino acid sequence alignment of Rv2170 and its orthologs present in pathogenic and non-pathogenic mycobacteria. Significant amino acid substitutions are highlighted.

**Supplementary Figure 6.**
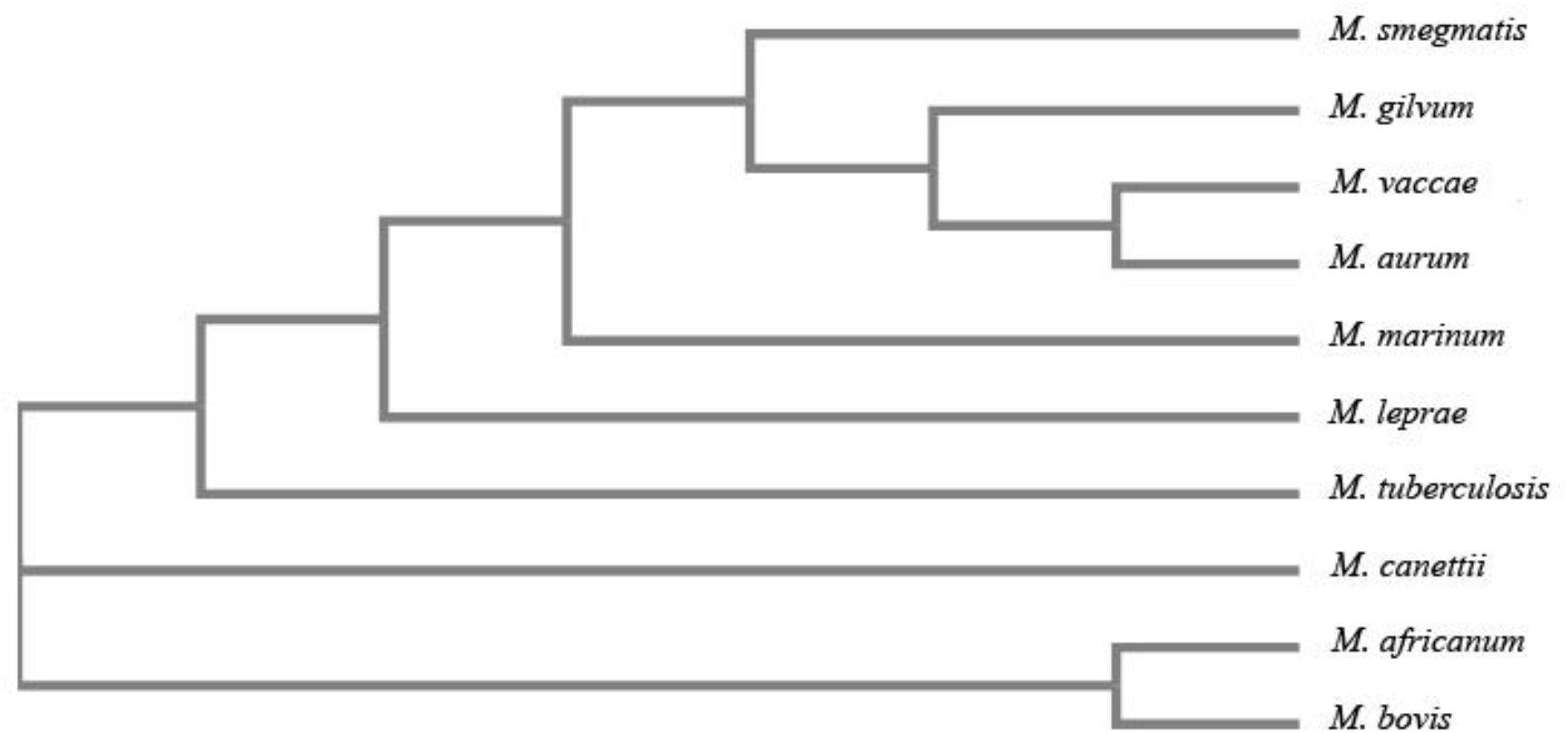
Cladogram of Rv2170 of *M. tuberculosis* H37Rv and its orthologs in pathogenic and non-pathogenic mycobacteria.

## References

1. Global tuberculosis report 2018. (2018) Geneva: World Health Organization; Licence: CC BY – NC-SA 3.0IGO

2. Mitchison, D. A. (2000) Role of individual drug in chemotherapy of tuberculosis. Int. J. Tuber. Lung. Dis. 4, 790–800

3. Timmins, G. S., and Deretic, V. (2006) Mechanisms of action of isoniazid. Mol. Microbiol. 62(5), 1220–1227

4. Ramaswamy, S., Reich, R., Dou, S., Jasperse, L., Pan, X., Wanger, A., Quitugua, T., and Graviss, E. A. (2003) Single nucleotide polymorphisms in genes associated with isoniazid resistance in Mycobacterium tuberculosis. Antimicrob. Agents Chemother. 47, 1241–1250

5. Silva, M. S., Senna, S. G., Ribeiro, M. O., Valim, A. R., Telles, M. A., Kritski, A., Morlock, G. P., Cooksey, R. C., Zaha, A., and Rossetti, M. L. (2003) Mutations in katG, inhA, and ahpC genes of Brazilian isoniazid-resistant isolates of Mycobacterium tuberculosis. J. Clinical Microbiol. 41(9), 4471–4474

6. Johnson, R., Streicher, E., Louw, G., Warren, R. M., van Helden, P. D., Victor, T. C. (2006) Drug resistance in *Mycobacterium tuberculosis*. Curr. Issues Mol. Biol. 8, 97–111

7. Dever, L. A., and Dermody, T. S. (1991) Mechanisms of bacterial resistance to antibiotics. Arch Intern Med. 151(5), 886–895

8. Wright, G. D. (2005) Bacterial resistance to antibiotics: Enzymatic degradation and modification. Adv. Drug Deliv. Rev. 57, 1451–1470

9. Munita, J. M., and Arias, C. A. (2016) Mechanisms of Antibiotic Resistance. Microbiol Spectr. 4(2), 10.1128/microbiolspec.VMBF-0016-2015

10. Jose, L., Ramachandran, R., Bhagavat, R., Gomez, R.L., Chandran, A., Raghunandanan, S., Omkumar, R.V., Chandra, N., Mundayoor, S., and Kumar, R.A. (2016) Hypothetical protein Rv3423.1 of *Mycobacterium tuberculosis* is a histone acetyltransferase. FEBS J. 283(2), 265–281

11. Favrot, L., Blanchard, J. S, and Vergnolle, O. (2016) Bacterial GCN5-related N-acetyltransferases: From resistance to regulation. Biochemistry 55(7), 989–1002

12. Mahapatra, S., Woolhiser, L. K., Lenaerts, A. J., Johson, J. L., Eisenach, K. D., Joloba, M. L., Boom, W. H., and Belisle, J. T. (2011). A novel metabolite of antituberculosis therapy demonstrates host activation of isoniazid and formation of the isoniazid-NAD^+^ adduct. Antimicrob. Agents Chemother. 56(1), 28–35

13. da Silva, J. L. Mesquita, A. R. C., and Ximenes, E. A. (2009) In vitro synergic effect of β-lapachone and isoniazid on the growth of *Mycobacterium fortuitum* and *Mycobacterium smegmatis*. Mem. Inst. Oswaldo Cruz. 104(4), 580–582

14. Umesiri, F. E., Lick, A., Fricke, C., Funk, J., and Nathaniel, T. (2015) Anti-tubercular activity of EDTA and household chemicals against *Mycobacterium smegmatis*, a surrogate for multi-drug resistant tuberculosis. Eur. Sci. J. 11(21), 133–149

15. Heinrichs, M. T., May, R. J., Heider, F., Reimers, T., B Sy, S. K., Peloquin, C. A., Derendorf, H. (2018) *Mycobacterium tuberculosis* strains H37ra and H37rv have equivalent minimum inhibitory concentrations to most antituberculosis drugs. Int J Mycobacteriol. 7, 156–61

16. Flentie, K., Harrison, G. A., Tukenmez, H., Livny, J., Good, J.A.D., Sarkar, S., Zhu, D. X., Kinsella, R. L., Weiss, L. A., Solomon, S. D., Schene, M. E., Hansen, M. R., Cairns, A. G., Kulen, M., Wixe, T., Lindgren, A. E. G., Chorell, E., Bengtsson, C., Krishnan, K. S., Hultgren, S. J., Larsson, C., Almqvist, F., and Stallings, C. L. (2019) Chemical disarming of isoniazid resistance in *Mycobacterium tuberculosis*. Proc. Natl. Acad. Sci. 116 (21), 10510–10517

17. Global tuberculosis report 2019. (2019) Geneva: World Health Organization; Licence: CC BY-NC-SA 3.0 IGO

18. Isakova, J., Nurmira, S., Denis, V., Zoy, G., Elnura, T., Nazira, A., and Almaz, A. (2018) Mutations of rpoB, katG, inhA and ahp genes in rifampicin and isoniazid-resistant *Mycobacterium tuberculosis* in Kyrgyz Republic. BMC Microbio. 18(1), 22

19. Wang, P., Pradhan, K., Zhong, X. B., Ma, X. (2016) Isoniazid metabolism and hepatotoxicity. Acta. Pharm. Sin. B. 6(5), 384–392

20. Guo, H., Seet, Q., Denkin, S., Parsons, L., and Zhnag, Y. (2006) Molecular characterization of isoniazid-resistant clinical isolates of *Mycobacterium tuberculosis* from the USA. J. Med. Microbio. 55, 1527–1531

21. Cardoso, R. F., Cardoso, M. A., Leite, C. Q. F., Sato, D. N., Mamizuka, E. M., Hirata, R. D. C., de Mello, F. F., and Hirata, M. H. (2007) Characterization of ndh gene of isoniazid resistant and susceptible Mycobacterium tuberculosis isolates from Brazil. Mem. Inst. Oswaldo Cruz. 102(1), 59–61

22. Torres, J. N., Paul, L. V., Rodwell, T. C., Victor, T. C., Amallraja, A. M., Elghraoui, A., Goodmanson, A. P., Ramirez-Busby, S. M., Chawla, A., Zadorozhny, V., Streicher, E. M., Sirgel, F. A., Catanzaro, D., Rodrigues, C., Gler, M. T., Crudu, V., Catanzaro, A., and Valafar, F. (2015) Novel *katG* mutations causing isoniazid resistance in clinical *M. tuberculosis* isolates. Emerg. Microbes Infect. 4(1), 1–9

23. Reygaert W. C. (2018) An overview of the antimicrobial resistance mechanisms of bacteria. AIMS Microbio. 4(3), 482–501

24. Schwarz, S., Kehrenberg, C., Doublet, B., and Cloeckaert, A. (2004) Molecular basis of bacterial resistance to chloramphenicol and florfenicol. FEMS Microbio. Rev. 28, 519–542

25. Huang, L., Yuan, H., Liu, M. F., Zhao, X. X., Wang, M. S., Jia, R. Y., Chen, S., Sun, K. F., Yang, Q., Wu, Y., Chen, X. Y., Cheng, A. C., and Zhu, D. K. (2017) Type B chloramphenicol acetyltransferases are responsible for chloramphenicol resistance in *Riemerella anatipestifer*, China. Front Microbiol. 8, 297

26. Houghton, J. L., Green, K. D., Pricer, R. E., Mayhoub, A. S., and Garneau-Tsodikova, S. (2013) Unexpected N-acetylation of capreomycin by mycobacterial Eis enzymes. J. Antimicrob. Chemother. 68(4), 800–805

27. Wang, X., Yang, S., Gu, J., and Deng, J. (2016) *Mycobacterium tuberculosis* arylamine *N*-acetyltransferase acetylates and thus inactivates *para*-aminosalicylicacid. Antimicrob. Agents Chemothera. 60(12), 7505–7508

28. Shi P, Zhang Y, Qu H., and Fan X. (2011) Systematic characterisation of secondary metabolites from Ixerissonchifolia by the combined use of HPLCTOFMS and HPLC-ITMS. Phytochem. Analy. 22, 66–73

29. Agrawal, P., Miryala, S., and Varshney, U. (2015) Use of *Mycobacterium smegmatis* deficient in ADP ribosyltransferaseas surrogate for *Mycobacterium tuberculosis* in drug testing and mutation analysis. PLoS ONE 10(4), e0122076

30. De Castro, E., Sigrist, C. J. A., Gattiker, A., Bulliard, V., Langendijk-Genevaux, P. S., Gasteiger, E., Bairoch, A., and Han ulo, N. (2006) ScanProsite: detection of PROSITE signature matches and ProRule-associated functional and structural residues in proteins. Nucleic Acids Res. 34, W362–365

31. Kuninger, D., Lundblad, J., Semirale, A., and Rotwein, P. (2007) A non-isotopic in vitro assay for histone acetylation. J. Biotechnol. 131(3), 253–260.

32. Martin, A., Camacho, M., Portaels, F., and Palomino, J. C. (2003) Resazurin microtiter assay plate testing of *Mycobacterium tuberculosis* susceptibilities to second-line drugs: rapid, simple, and inexpensive method. Antimicrob. Agents Chemother. 47(11), 3616–3619

33. Altschul, S. F., Gish, W., Miller, W., Myers, E. W., Lipman D. J. (1990) Basic local alignment search tool. J. Mol. Bio. 215(3), 403–410

34. Fiser, A., and Sali, A. (2003) Modeller: generation and refinement of homology-based protein structure models. Methods Enzymol. 374, 461–491

35. Laskowski, R. A., MacArthir, M. W., Moss, D. S. Thornoton, J. M. (1993) PROCHECK: a program to check the stereochemical quality of protein structures. J. Appl. Crystaallogr. 26(2), 283–291

36. Binkowski, T. A., Naghibzadeh, S., and Liang, J. (2003). CASTp: computed atlas of surface topography of proteins. Nucleic Acids Res. 31(13), 3352–3355

37. Morris, G. M., Huey, R., Lindstorm, W., Sanner, M. F., Belew, R. K., Goodsell, D. S., Olson, A. J. (2009) AutoDock4 and AutoDockTools4: Automated docking with selective receptor flexibility. J. Comput. Chem. 30(16), 2785–2791

